# Long wavelength light reduces the negative consequences of dim light at night in the *Cntnap2* mouse model of autism

**DOI:** 10.1101/2022.06.09.494760

**Authors:** Huei Bin Wang, David Zhou, Shu Hon Christopher Luk, Hye In Cha, Amanda Mac, Rim Chae, Anna Matynia, Ben Harrison, Sina Afshari, Gene D. Block, Cristina A. Ghiani, Christopher S. Colwell

**Affiliations:** Molecular, Cellular, Integrative Physiology Graduate Program, David Geffen School of Medicine; University of California Los Angeles; Department of Psychiatry & Biobehavioral Sciences, David Geffen School of Medicine; University of California Los Angeles; Laboratory of Ocular Molecular and Cellular Biology and Genetics, Jules Stein Eye Institute, David Geffen School of Medicine; University of California Los Angeles; Department of Pathology and Laboratory Medicine, David Geffen School of Medicine; University of California Los Angeles; Intellectual and Developmental Disabilities Center, David Geffen School of Medicine; University of California Los Angeles; Korrus Inc Los Angeles, CA

**Keywords:** Autism spectrum disorder, amygdala, c-Fos, circadian, *Cntnap2* knock out, light pollution, melanopsin, mice

## Abstract

Many patients with autism spectrum disorders (ASD) show disturbances in their sleep/wake cycles, and may be particularly vulnerable to the impact of circadian disruptors. We have previously shown that exposure to dim light at night (DLaN) in contactin associated protein-like 2 knock out (*Cntnap2* KO) mice disrupts diurnal rhythms, increases repetitive behaviors while reducing social interactions. These negative effects of DLaN may be mediated by intrinsically photosensitive retinal ganglion cells (ipRGCs) expressing the photopigment melanopsin, which is maximally sensitive to blue light (480nm). In this study, we used a light-emitting diode (LED) array that enabled us to shift the spectral properties of the DLaN while keeping the intensity at 10 lx. First, using wild-type (WT) mice, we confirmed that the short-wavelength enriched lighting produced strong acute suppression of locomotor activity (masking), robust light-induced phase shifts, and c-Fos expression in the suprachiasmatic nucleus, while the long-wavelength enriched lighting evoked much weaker responses. Furthermore, exposure of WT mice to the short-wavelength light at night reduced the amplitude of locomotor activity rhythms and impaired social interactions. Mice lacking the melanopsin expressing ipRGCs (*Opn4*^DTA^ mice) were resistant to these negative effects of DLaN. Importantly, the shift of the DLaN stimulus to longer wavelengths ameliorated the negative impact on the activity rhythms and autistic behaviors (i.e. reciprocal social interactions, repetitive grooming) of the *Cntnap2* KO model. The short-, but not the long-wavelength enriched, DLaN triggered cFos expression in the peri-habenula region as well as in the basolateral amygdala (BLA). Finally, DLaN-driven c-Fos induction in BLA glutamatergic neurons was about 3-fold higher in the *Cntnap2* KO mice, suggesting that these cells may be particularly vulnerable to the effects of photic disruption. Broadly, our findings suggest that the spectral properties of light at night should be considered in the management of ASD and other neurodevelopmental disorders.

## Introduction

Dim light at night (DLaN) is a common environmental perturbation of the circadian timing system and has been linked to a range of negative consequences (Stevenson et al., 2015; Lunn et al., 2017). Prior work in pre-clinical models has demonstrated that light at night negatively impacts the metabolism (Fonken et al., 2013a; Plano et al., 2017), immune function (Bedrosian et al., 2011; Fonken et al., 2013b; Lucassen et al., 2016), mood and cognition (Lazzerini Ospri et al., 2017; An et al., 2020; Walker et al., 2020). Individuals diagnosed with autism spectrum disorders (ASD) commonly experience disturbances in their circadian rhythms, and commonly find difficulty in both falling asleep as well as sleeping through the night (Cohen et al., 2014; Devnani and Hegde, 2015; Mazurek and Sohl, 2016; Baker and Richdale, 2017; Shelton and Malow, 2021). Perhaps because of this difficulty sleeping at night, ASD patients spend more time exposed to the electronic screens at night compared with age-matched controls (Engelhardt et al., 2013; Stiller et al., 2019; Healy et al., 2020; Dong et al., 2021). It has been speculated that this exposure to DLaN via screens and other lighting sources could be detrimental to ASD populations. In support of this speculation, we previously reported that an animal model with a genetic risk factor associated with ASD (contactin associated protein-like 2, *Cntnap2*) shows reduced social preference and increased stereotypic grooming behavior after a 2-week exposure to DLaN (Wang et al., 2020).

The negative effects of nightly light exposure could be mediated by the intrinsically photosensitive retinal ganglion cells (ipRGCs) expressing the photopigment melanopsin, which is maximally sensitive to shorter wavelengths (λ) of light with a peak response to 480nm light (Hattar et al., 2002; Panda et al., 2005). In addition, ipRGCs receive inputs from rod and cone photoreceptors (Hattar et al., 2002; Lucas et al., 2012; Van Diepen et al., 2021) with the rods driving the circadian response to sensitivity to low-intensity light (Altimus et al., 2010; Lall et al 2010). These findings indicate that the circadian system is sensitive to broad spectrum of light (Foster et al., 2020), and raise the question of whether changes in the spectral properties of DLaN matter. Perhaps the intensity of the night-time illumination is the only parameter that is biologically important? In fact, recent studies are beginning to address the issue of whether adjusting the spectral composition of light to reduce melanopic stimulation can be used as a strategy to minimize disruption of circadian rhythms (Gladanac et al., 2019; Nagare et al., 2019; Figueiro and Pedler, 2020; Mouland et al., 2021; Vethe et al., 2021).

In the present study, we employed an array of light-emitting diodes (LEDs) to generate dim illumination (10 lx) with a spectral composition that was enriched in either shorter wavelengths (λ) designed to activate melanopsin or longer λ designed to minimize activation of this photopigment. First, we ascertained in wild-type (WT) mice whether our two light treatments differentially regulated negative masking behavior, light-induced phase shifts of circadian behavior as well as the light-evoked c-Fos response in the suprachiasmatic nucleus (SCN). A transgenic line of mice (*Opn4*^DTA^) lacking melanopsin and with diminished ipRGCs due to the targeted expression of diphtheria toxin A in the *Opn4* expressing cells (Matynia et al., 2012; Chew et al., 2017) was used to further evaluate the contribution of the ipRGCs in mediating the effects of DLaN. Next, we determined whether the two light treatments differentially impacted DLaN-driven changes in daily patterns of activity, sociability, and grooming in WT and *Cntnap2* knock-out (KO) mice. Finally, we examined whether short-λ enhanced DLaN altered light-evoked changes in the number of c-Fos+ cells in selected brain regions in both WT and *Cntnap2* KO mice.

## Materials and Methods

### Animals

All animal procedures were performed in accordance with the UCLA animal care committee’s regulations. Both adult male and female mice (3-4 months old, mo) were used in this study. *Cntnap*^*2tm2Pel*^*/*J mutant mice backcrossed to the C57BL/6J background strain (B6.129(Cg)-*Cntnap2*^*tm2Pele*^/J, stock number 017482; RRID:IMSR_JAX:017482) and the control C57BL/6J strain (stock number 000664; RRID:IMSR_JAX:000664) were from our breeding colony maintained in an approved facility of the Division of Laboratory Animal Medicine at the University of California, Los Angeles (UCLA). *Cntnap2* KO and WT mice were obtained from heterozygous breeding pairs, weaned at postnatal day 21 and genotyped (TransnetYX, Cordova, TN). *Opn4*^DTA^ mice (*Opn4*^tm3.1(DTA)Saha^/J; stock number 035927; RRID:IMSR_JAX:035927) and their WT littermates were kindly provided by Dr. Anna Matynia (Jules Stein Eye Institute, UCLA). Vglut2-ires-Cre knock-in mice (B6J.129S6(FVB)-Slc17a6^tm2(cre)Lowl^/MwarJ; stock number 028863; RRID:IMSR_JAX:028863) carrying a Cre recombinase targeting excitatory glutamatergic neurons, as well as Ai6 mice (B6.Cg-Gt(ROSA)26Sor^tm6(CAG-ZsGreen1)Hze^/J; stock number 007906; RRID:IMSR_JAX:007906) expressing robust ZsGreen1 fluorescence following Cre-mediated recombination were obtained from Jackson Laboratory. The Ai6 is a Cre reporter allele designed to have a loxP-flanked STOP cassette preventing transcription of a CAG promoter-driven enhanced green fluorescent protein variant (ZsGreen1) -all inserted into the Gt(ROSA)26Sor locus. We bred this floxed reporter gene into the Cntnap2 mutant to create *Cntnap2* KO mice heterozygous for Vglut2 Cre-Ai6 (ZsGreen1). Cohorts of male and female mice were group housed prior to experimentation.

### Lighting manipulations

Mice were housed in light-tight ventilated cabinets in temperature- and humidity-controlled conditions, with free access to food and water. All the mice were first entrained to a normal lighting cycle: 12:12 hr light:dark (LD). Light intensity during the day was 300 lux as measured at the base of the cage, and 0 lux during the night. The time of lights-on defines Zeitgeber Time (ZT) 0 and the time of lights-off defines ZT 12. Following entrainment to normal LD, mice were singly housed under different lighting conditions for 2 weeks: LD, night light with the spectral composition aimed at minimizing the stimulation of melanopsin (long λ enriched lights), or light with the spectral composition aimed at maximizing the stimulation of melanopsin (short λ-enriched lights). The treated mice were exposed nightly to one of these lights from ZT 12 to 24. Control mice were held on the normal LD 12:12 schedule. The short- and long wavelength (10 lx intensity) were generated by the Korrus Inc (Los Angeles, CA) LED lighting system (**Fig. 1A)**. Following recent recommendations (Lucas et al., 2014), the LED output was measured using UPRtek spectrophotometer (Model MK350S, Taiwan) at the level of the cage floor. Using the International Commission on Illumination (CIE) S 026 α-opic Toolbox (DOI: 10.25039/S026.2018), the melanopic α-opic irradiance of the long λ enhanced output was estimated at 0.001 W/m^2^ while short λ enhanced output was estimated to be 0.020 W/m^2^. The melanopic α-opic equivalent daylight (D65) illuminance of the long λ output was estimated at 1 melanopic lx while the short λ output was estimated at 15 melanopic lx. The total energy of the long λ enhanced light was measured as an illumination of 10.4 lx, irradiance of 0.05 W/m^2^, log photon irradiance of 17.2 s^1^/m^2^ while the short λ enhanced light was measured as an illumination of 10.0 lx, irradiance of 0.03 W/m^2^, log photon irradiance of 17.0 s^1^/m^2^.

**Fig. 1:**
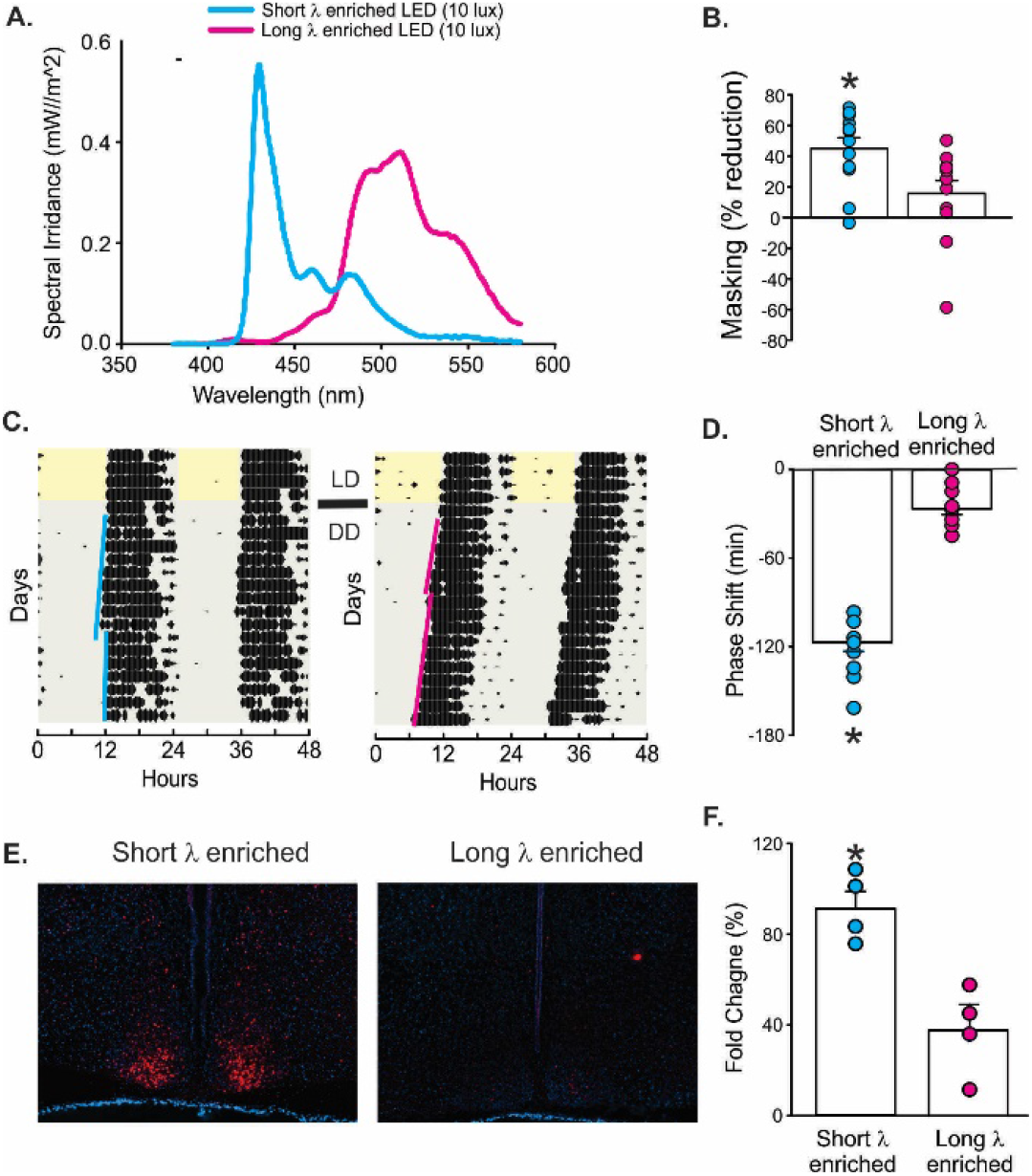
Shift to longer wavelength illumination reduces negative masking, light-induced phase shifts of the circadian activity rhythms, and induction of c-Fos in the suprachiasmatic nucleus of WT mice. **(A)** Spectral irradiance of the light emitted by the LED array using two lighting treatments: a short wavelength (λ) enriched (10 lx, peak at 478 nm) and a long λ enriched illumination (10 lx, peak at 608 nm). **(B)** Light-induced masking of activity in WT mice exposed to either the short λ or long λ enriched light for one hour at ZT 14. The activity level during this light exposure was compared to that during the equivalent hour (ZT 14-15) on the day before the treatment (baseline activity). **(C)** Examples of light-induced phase shifts of wheel-running activity onsets. After entrainment to the standard light:dark (LD) cycle, WT mice were placed into constant darkness (DD) for 10 days and then exposed to either short λ or long λ enriched illumination (10 lx, 15 min) at CT 16 (4 hours after the activity onset). Solid lines show lines running through activity onset before and after light treatment. **(D)** The short λ enhanced illumination produced significant phase delays of the circadian system. **(E)** Examples of light-evoked c-Fos expression in the suprachiasmatic nucleus (SCN). Mice were exposed to either the short λ or long λ enriched illumination (10 lx) for 15 min at CT 16, perfused 45 min later and the brains collected. (**F**) The short λ enhanced illumination produced a significant increase in the number of c-Fos positive cells in the SCN. Cells were counted in both the left and right SCN in two consecutive coronal sections per animal and the numbers averaged to obtain one value per animal. The histograms show the means ± the standard error of the mean (SEM). Data were analyzed using one-way ANOVA (**Table 1**) followed by the Holm-Sidak multiple comparisons test with the asterisks indicating significant difference between the groups (*P* < 0.05).

Daily pattern of activity was monitored using a top-mounted passive infra-red (IR) motion detector reporting to a VitalView data recording system (Mini Mitter, Bend, OR) as previously described (Wang et al., 2017; 2018). Detected movements were recorded in 3 min bins, and 10 days of data were averaged for analysis using the El Temps chronobiology program (A. Diez-Noguera, Barcelona, Spain; http://www.el-temps.com/principal.html). The strength of the rhythm was determined from the amplitude of the χ2 periodogram at 24 hr, to produce the rhythm power (%V) normalized to the number of data points examined. The periodogram analysis uses a χ2 test with a threshold of 0.001 significance, from which the amplitude of the periodicities is determined at the circadian harmonic to obtain the rhythm power. The percentage of sleep-phase activity was determined by the relative distribution of activity during the day versus the night. Fragmentation and onset variability were determined using the Clocklab program (Actimetrics, Wilmette, IL; http://actimetrics.com/products/clocklab/). Fragmentation was determined by the number of activity bouts. Each bout was counted when activity bouts were separated by a gap of 21 min. The onset variability was by first determining the daily onset of activity over 10 days, and then determining the deviation from the best-fit line drawn through the 10 onset times. To produce the representative waveforms, hourly running averages of the same series of activity data was plotted against time of day.

**Table 1.** Reduced circadian outputs resulted from the long wavelength illumination. WT mice were used to test whether the long wavelength light has a diminished impact on 3 traditional assays for light input to the circadian system. For masking, mice were exposed to light (10 lx, 60 min) from ZT 14 to 15. For phase shifts, mice were held in constant darkness (DD) and exposed to light at CT 16 (10 lx, 15 min). For c-Fos expression, mice held in DD were exposed to light at CT 16 (10 lx, 15min), euthanized at CT 17 and the brains collected. The number of immunopositive cells in both the left and right SCN from two consecutive coronal sections per animal were averaged to obtain one number per animal. An equal number of male and female mice was used, and values are shown as the mean ± SEM. Data from the behavioral tests passed the normality and equal variance tests and differences were evaluated by one way ANOVA followed by the Holm-Sidak Multiple comparison method. The c-Fos expression data did not past the equal variance test and the Kruskal-Wallis one way ANOVA on ranks was used to evaluate significances. In this and subsequent tables, *P* ≤ 0.05 was considered significant, and significant differences are shown in bold.

| Test | WT untreated | Short $\lambda$ enriched | Long $\lambda$ enriched | One way ANOVA |
| --- | --- | --- | --- | --- |
| Masking (% reduction) | -3.0 ± 6.9 (n=14) | -51.0 ± 5.0 (n=11) | -13.6 ± 7.7 (n=11) | **$F_{(2, 35)}=13.7$; $P<0.001$** |
| Phase shift (min) | -3.7 ± 4.0 (n=8) | -117.2 ± 5.9 (n=12) | -26.4 ± 3.9 (n=12) | **$F_{(2, 31)}=149.3$; $P<0.001$** |
| c-Fos+ in SCN | 5.6 ± 0.4 (n=4) | 94.2 ± 6.5 (n=4) | 18.8 ± 2.3 (n=4) | **$H_{(2)}=2.651$; $P<0.001$** |

### Photic suppression on nocturnal activity (negative light masking)

A separate cohort of WT mice (3-4 mo) were housed under a 12:12 LD cycle to determine the effects of LED lights (Korrus Inc) at night on masking of nocturnal activity. Animals were individually housed in cages with IR motion detector in light-tight chambers with the same housing conditions as described above. Their locomotor activity was recorded per 3-min interval and monitored to ensure the entrainment to the housing LD cycles. The animals were subjected to one hour of light (enriched for long or short wavelengths) at ZT 14. The activity level during this light exposure was compared to the activity level during the equivalent hour (ZT 14 to 15) on the day before the treatment. The fold change (%) was reported.

### Phase shift of activity rhythms

After the habituation to running wheel cages, a different cohort of WT mice (3-4 mo) were released into constant darkness (DD) for 10 days. The circadian time (CT) of their free-running sleep/wake cycles were monitored with the VitalView. The time of activity onset under DD was defined as CT 12. On day 11, mice were exposed to long-or short-wavelength light (10 lx, 60 min) at CT 16. After the treatment, the mice stayed in DD for following 10 days. And best-fit lines of the activity onsets of sleep/wake cycles before and after the treatment were determined and compared.

### Photic induction of c-Fos in the SCN and Immunofluorescence

A separate cohort of WT mice (n= 2 male and 2 female, 3-4 mo) was housed in DD conditions. The animals were exposed to long-or short-wavelength light (10 lx) for 15 mins at CT 16. Forty-five mins later, the mice were euthanized with isoflurane (30%–32%) and transcardially perfused with phosphate-buffered saline (PBS, 0.1 M, pH 7.4) containing 4% (w/v) paraformaldehyde (Sigma). The brains were rapidly dissected out, post-fixed overnight in 4% PFA at 4 °C, and cryoprotected in 15% sucrose. Sequential coronal sections (50 μm), containing the middle SCN, were collected using a cryostat (Leica, Buffalo Grove, IL) and then further processed for c-Fos immunofluorescence as reported below (Wang et al., 2017).

### Behavioral tests

The reciprocal social interaction test was used to assess general sociability and interest in social novelty. On days of testing, the animals were taken out from the light-tight chambers, and habituated to the testing room for at least 30 mins. To avoid sleep disruptions, all behavioral tests were conducted in the middle of the dark phase (ZT 16-18). The social behavior was evaluated by familiarizing the testing mouse with the testing arena first (habituation for 30 mins), and then introducing a never-met stranger mouse into the same arena. The stranger mouse had the same age, sex, and genotype as the testing mouse. The two mice were allowed to interact for 10 mins, and the testing trials were recorded. Duration of sniffing (nose to nose sniff and nose to anogenital sniff), social grooming (the testing mouse grooms the novel stranger mouse on any part of the body) and touching was timed and reported for the time of social behavior.

The grooming test were performed to evaluate the level of their stereotypic behavior (ZT 16-18). After the habituation to the testing room for at least 30 mins, the testing mice were introduced to the testing arena and their behavior was recorded by a camcorder for 30 mins. The duration of grooming behavior was scored as previously described (Wang et al., 2020).

### Immunofluorescence

At the end of the 2 weeks of exposure to the 3 different lighting conditions, mice were anesthetized with isoflurane at a specific time during the night (ZT 18) and transcardially perfused with phosphate buffered saline (PBS, 0.1 M, pH 7.4) containing 4% (w/v) paraformaldehyde (Sigma). The brains were rapidly dissected out, post-fixed overnight in 4% PFA at 4 °C, and cryoprotected in 15% sucrose. Coronal sections (50μm) were obtained using a cryostat (Leica, Buffalo Grove, IL), collected sequentially, and paired along the anterior– posterior axis before further processing. Immunohistochemistry was performed as previously described (Wang et al., 2017; Lee et al., 2018). Briefly, free-floating coronal sections containing the brain regions of interest were blocked for 1 hr at room temperature (1% BSA, 0.3% Triton X-100, 10% normal donkey serum in 1xPBS) and then incubated overnight at 4°C with a rabbit polyclonal antiserum against cFos (1:1000, Millipore and Cell Signaling) followed by a Cy3-conjugated donkey-anti-rabbit secondary antibody (Jackson ImmunoResearch Laboratories, Bar Harbor, ME). Sections were mounted and coverslips were applied with Vectashield mounting medium containing DAPI (4’-6-diamidino-2-phenylinodole; Vector Laboratories, Burlingame, CA). Sections were visualized on a Zeiss AxioImager M2 microscope (Zeiss, Thornwood NY) equipped with an AxioCam MRm and the ApoTome imaging system

### c-Fos positive Cell Counting

Images for counting were acquired using either a 20X or 40X objective and the Zeiss Zen digital imaging software. Both male and female mice, in equal number, were used for each assessment and two observers masked to genotype and experimental group performed the cell countings.

#### Photic induction of c-Fos in the SCN

Images of both the left and right middle SCN were acquired with a 20X objective using the tile tool of the Zen software. The SCN was then manually traced in each image using the NIH ImageJ software (https://imagej.net/software/fiji/), and the cells immunopositive for c-Fos were counted with the aid of the cell counter plugin of ImageJ in two consecutive sections at the level of the mid-SCN. Values from the left and right SCN in the two slices were averaged to obtain one value per animal and are presented as the mean ± SEM of 4 animals/experimental group.

#### c-Fos expression in different brain regions of interest

Images of all the regions of interest (both right and left hemisphere) were acquired with a 20X objective using the tile tool of the Zen software. Each brain area was manually traced to define the regions of interest and all the cells immunopositive for cFos were counted with the aid of the cell counter plugin of ImageJ in two consecutive sections. Values from the left and right hemispheres in the two slices were averaged to obtain one value per animal and are presented as the mean ± SEM of 6 animals/genotype/experimental group.

#### c-Fos expression in the Basolateral Amygdala of vGlut2-ZsGreen1 expressing mice

Seven to fifteen images from the left and right Basolateral Amygdala (BLA) were randomly acquired with a 40X objective from 6 consecutive slices/animal. Two investigators masked to genotype and treatment group performed the cell counting using the Zen software tools in 5-7 images/animal. The followings were determined in each image: (1) the total number of vGlut-ZsGreen1 cells; (2) the number of vGlut-ZsGreen1 cells immunopositive for c-Fos. The percentage of cFos positive cells/vGlut-ZsGreen1 cells per image was determined and then averaged to obtain a single value/animal. On average, each image contained about 30-45 vGlut-ZsGreen1 cells, with the *Cntnap2* KO exposed to short λ enriched DLaN had the lowest counts with an average of 33 cells, while the other three groups had on average about 45 cells. Data are shown as the mean ± SEM of 4 animals/genotype/ experimental group, 2 males and 2 females were present in each group.

### Statistics

SigmaPlot (version 12.5, SYSTAT Software, San Jose, CA) was used to run statistical analyses. Cohorts of mice kept in LD are referred to as baseline. One-way analysis of variance (ANOVA) was used to determine the significance of the impact of long or short wavelength enriched DLaN on the three assays for the light input to the circadian system in WT mice (**Table 1**). Given the low sample size of the *Opn4*^DTA^ mice and littermates, we used two-tailed *t*-test or the Mann Whitney rank-sum test to determine whether DLaN significantly altered the mice activity parameters (**Table 2**). A two-way ANOVA was used to analyse the activity rhythms with time and treatment as factors, as well as the behavioral tests (**Table 3**) and the number of c-Fos^+^ cells (**Table 4**) with genotype and treatment as factors. The Holm-Sidak test for multiple comparisons was used when appropriate. Values are reported as the mean ± standard error of the mean (SEM). Differences were determined significant if *P* < 0.05.

**Table 2.**
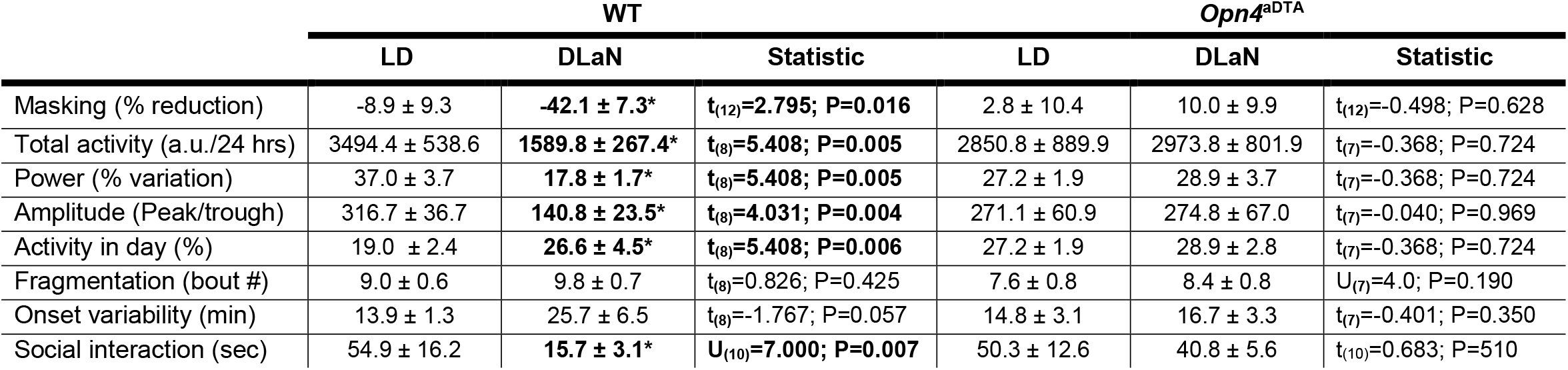
*Opn4*^*DTA*^ mice were less affected by short wavelength enriched DLaN. Light input to the circadian system is mediated by the ipRGC population that can be selectively ablated in a line of mice with the gene coding for melanopsin (OPN) expressing a component of the diphtheria toxin. We compared the impact of short wavelength enriched DLaN on the activity rhythms and social interaction behavioral test in WT and *Opn4*^DTA^ littermates. Both male and female mice were used. Values are shown as means ± SEM of n=4-6 animals/genotype/light condition for the activity rhythms and n=6-8 animals/genotype for the social interaction behavioral test. Because of the sample size, the effects of the short wavelength enriched DLaN was analyzed only within the genotype, and data were analyzed with a two-tailed *t*-test. For parameters that did not pass normality tests, the Mann Whitney rank-sum test was used and the U values reported. P values < 0.05 were considered significant. The asterisks indicate significant differences between LD and DLaN treatments within genotype. Significant differences are shown in bold. a.u.= arbitrary units

**Table 3.** Long wavelength enriched illumination minimized DLaN disruption. Comparisons of age-matched WT and *Cntnap2* KO mice under LD or DLaN regimen (n = 12-15/group). Values are shown as averages ± SEM. Data were analyzed with a two-way ANOVA using genotype and treatment as factors. The Holm-Sidak test for multiple comparisons was used when appropriate. The asterisks indicate significant differences between LD and DLaN treatments within genotype and the crosshatches between genotypes. Significant differences are shown in bold. a.u.= arbitrary units

|  | WT |  |  | *Cntnap2* KO |  |  | Two-way ANOVA |  |
| --- | --- | --- | --- | --- | --- | --- | --- | --- |
| | Baseline LD | Short $\lambda$ DLaN | Long $\lambda$ DLaN | Baseline LD | Short $\lambda$ DLaN | Long $\lambda$ DLaN | Genotype | Treatment |
| Total activity (a.u./24 hrs) | 3003.9 ± 321 | 2114.3 ± 237 | 3443.3 ± 524 | **1943.4±135<sup>#</sup>** | 1639.4 ± 254 | 2145.6 ± 217 | **F<sub>(1,95)</sub>=14.16; P<0.001** | **F<sub>(2,95)</sub>=3.88; P=0.024** |
| Power (% variance) | 31.8 ± 1.1 | **22.6 ± 1.0\*** | 30.6 ± 2.2 | **24.8 ± 1.2<sup>#</sup>** | **19.3 ± 1.4\*** | 28.9 ± 2.0 | **F<sub>(1,95)</sub>=5.39; P=0.022** | **F<sub>(2,95)</sub>=10.97; P<0.001** |
| Amplitude (Peak/trough) | 250.1 ± 29.6 | **148.7 ± 1\*** | 295.3 ± 45.9 | **183.9 ± 15.9<sup>#</sup>** | **132.1 ± 19.9\*** | 212.1 ± 22.5 | **F<sub>(1,95)</sub>=5.49; P=0.018** | **F<sub>(2,95)</sub>=7.23; P=0.001** |
| Activity in day (%) | 18.7 ± 0.7 | **26.0 ± 1.9\*** | 19.6 ± 1.7 | 26.0 ± 2.0 | 32.3 ± 2.0 | 19.3 ± 1.6 | F<sub>(1,95)</sub>=2.80; P=0.098 | **F<sub>(2,95)</sub>=27.63; P<0.001** |
| Onset variability (min) | 18.2 ± 1.9 | **33.44 ± 5\*** | 25.0 ± 3.0 | 23.5 ± 2.0 | **42.0 ± 6.1\*** | 36.6 ± 2.9 | **F<sub>(1,95)</sub>=10.27; P=0.002** | **F<sub>(2,95)</sub>=16.10; P<0.001** |
| Fragmentation (# bouts) | 10.5 ± 0.9 | 10.7 ± 1.0 | 10.8 ± 2.0 | 11.6 ± 1.3 | 12.4 ± 1.5 | 11.6 ± 1.6 | F<sub>(1,95)</sub>=1.13; P=0.29 | F<sub>(2,95)</sub>=0.07; P=0.93 |
| Social interaction (sec) | 48.8 ± 6.2 | **26.3 ± 6.8\*** | 43.3 ± 9.6 | **23.4 ± 3.4<sup>#</sup>** | **8.6 ± 2.0\*** | 16.9 ± 3.9 | **F<sub>(1,95)</sub>=15.04; P<0.001** | **F<sub>(2,95)</sub>=4.54; P=0.013** |
| Grooming (sec) | 13.4 ± 1.1 | 15.7 ± 2.0 | 10.1 ± 1.6 | **31.2 ± 1.8<sup>#</sup>** | **43.7 ± 2.8\*** | 27.7 ± 2.5 | **F<sub>(1,151)</sub>=149.14; P<0.001** | **F<sub>(2,151)</sub>=11.51; P<0.001** |

**Table 4:** Sex differences in WT and *Cntnap2* KO mice held in LD conditions. Comparisons of age-matched male and female WT and *Cntnap2* KO mice held in LD. Values are shown as averages ± SEM. Data were analyzed with a two-way ANOVA using genotype and treatment as factors. The Holm-Sidak test for multiple comparisons was used when appropriate. There were no significant interactions between genotype and sex, thus these values are not reported. The crosshatches indicate significant differences between genotypes. Significant differences are shown in bold. a.u.= arbitrary units

| Measures | WT |  | *Cntnap2* KO |  | Two-way ANOVA |  |
| --- | --- | --- | --- | --- | --- | --- |
|  | Male<br>Baseline LD | Female<br>Baseline LD | Male<br>Baseline LD | Female<br>Baseline LD | Genotype | Sex |
|  | n=12 | n=11 | n=12 | n=12 |  |  |
| Total activity<br>(a.u./24 hrs) | 2756.8 ± 327.6 | 3276.6 ± 429.8 | 2073.8 ± 213.8 | **1812.9 ± 156.6<sup>#</sup>** | **$F_{(1,46)}=13.28; P<0.001$** | $F_{(1,46)} = 0.19; P=0.662$ |
| Power (% variance) | 31.9 ± 1.5 | 32.5 ± 1.5 | **24.5 ± 1.4<sup>#</sup>** | **24.3 ± 1.7<sup>#</sup>** | **$F_{(1,46)}=19.92; P<0.001$** | $F_{(1,46)} = 0.03; P=0.873$ |
| Amplitude<br>(Peak/trough) | 237.6 ± 29.2 | 251.4 ± 29.3 | 183.8 ± 14.9 | **167.2 ± 18.2<sup>#</sup>** | **$F_{(1,46)}=8.54; P=0.006$** | $F_{(1,46)} = 0.01; P=0.951$ |
| Social interaction<br>(sec) | 41.2 ± 6.4 | 56.0 ± 8.3 | 22.5 ± 5.2 | 22.5 ± 6.4 | **$F_{(1,47)}=15.62; P < 0.001$** | $F_{(1,47)}=1.26; P=0.268$ |
| Grooming (sec) | 14.4 ± 1.6 | 12.3 ± 1.6 | **35.8 ± 2.6<sup>#</sup>** | **28.3 ± 2.0<sup>#</sup>** | **$F_{(1,69)}=82.34; P<0.001$** | **$F_{(1,69)}=5.37; P=0.024$** |

## Results

### Long wavelength enriched illumination reduces negative masking, light-induced phase shifts of the circadian activity rhythm, and induction of c-Fos in the SCN of WT mice

In these first set of experiments, we evaluated the influence of long and short λ enhanced lighting (**Fig. 1A**) on three classic tests of the ipRGC input to the circadian system in WT mice. The effects on light-driven masking of activity were measured with a passive infra-red activity assay. While the short wavelength enriched lights (10 lx, 60 min, ZT 14) elicited a significant reduction (51 ± 5%) in masking behavior in the WT, the long wavelength illumination produced a low level of masking (14 ± 7%) (**Fig. 1B, Table 1)**. Next, we measured the impact of the two treatments on the light-induced phase shift of the circadian system using wheel-running activity (**Fig. 1C**). The WT mice were exposed to short-or long-wavelength lighting (10 lx, 15 min, CT 16) while activity was monitored in DD. The short-wavelength light triggered a significantly larger phase delay (−117 ± 6 min) in the mice compared to the long-λ (−26 ± 4 min) (**Fig. 1D, Table 1)**. Finally, we examined how the two lighting treatment (10 lx, 15 min, CT 16) affected the number of c-Fos immunopositive cells in the SCN (**Fig. 1E)**. The SCN of the mice exposed to the shorter wavelength light exhibited a robust induction of c-Fos (94 ± 6 cells per SCN section) whilst those exposed to the longer wavelength light had significantly fewer cells (19 ± 2 per SCN section) (**Fig. 1F, Table 1**). Hence, shifting the spectral properties of light toward longer wavelength significantly reduced the effect of light on negative masking, light-induced phase shifts and light-induction of c-Fos in the SCN.

### Melanopsin (Opn4) expressing cells mediate the influence of DLaN on both activity cycles and social behavior

It is well-established that the *Opn4* expressing ipRGCs mediate the effects of light on the circadian system and we sought to test their role in mediating the behavioral impact of DLaN. *Opn4*^DTA^ mice and their WT littermates were held under either LD control environment or the short λ enriched DLaN for 2 weeks. While the WT mice exhibited a robust reduction in the strength of the activity rhythms (activity levels, power, amplitude) when exposed to the short λ enriched DLaN, these parameters were unaltered in the mutant mice (**Fig. 2A, B, Table 2**). **N**ext, we examined negative masking behavior as described above. Again, the WT mice exhibited a drastic suppression of activity (42 ± 7%), while the *Opn4*^DTA^ did not show any significant light-evoked change on their activity (9 ± 9%; Fig. 2C, **Table 2**). We had previously shown that DLaN reduces social behavior in C57 mice (Wang et al., 2020), hence, the effect of the short λ illumination was examined on this behavior in the *Opn4*^DTA^. In WT mice, the DLaN greatly reduced social interactions (57.4 ± 10%), while the mutant mice were resistant to this negative effect of night-time illumination (8.9 ± 8.2%, **Fig. 2D, Table 2**). Together these data indicate that the ipRGCs are required for at least some of the negative effects of DLaN.

**Fig. 2:**
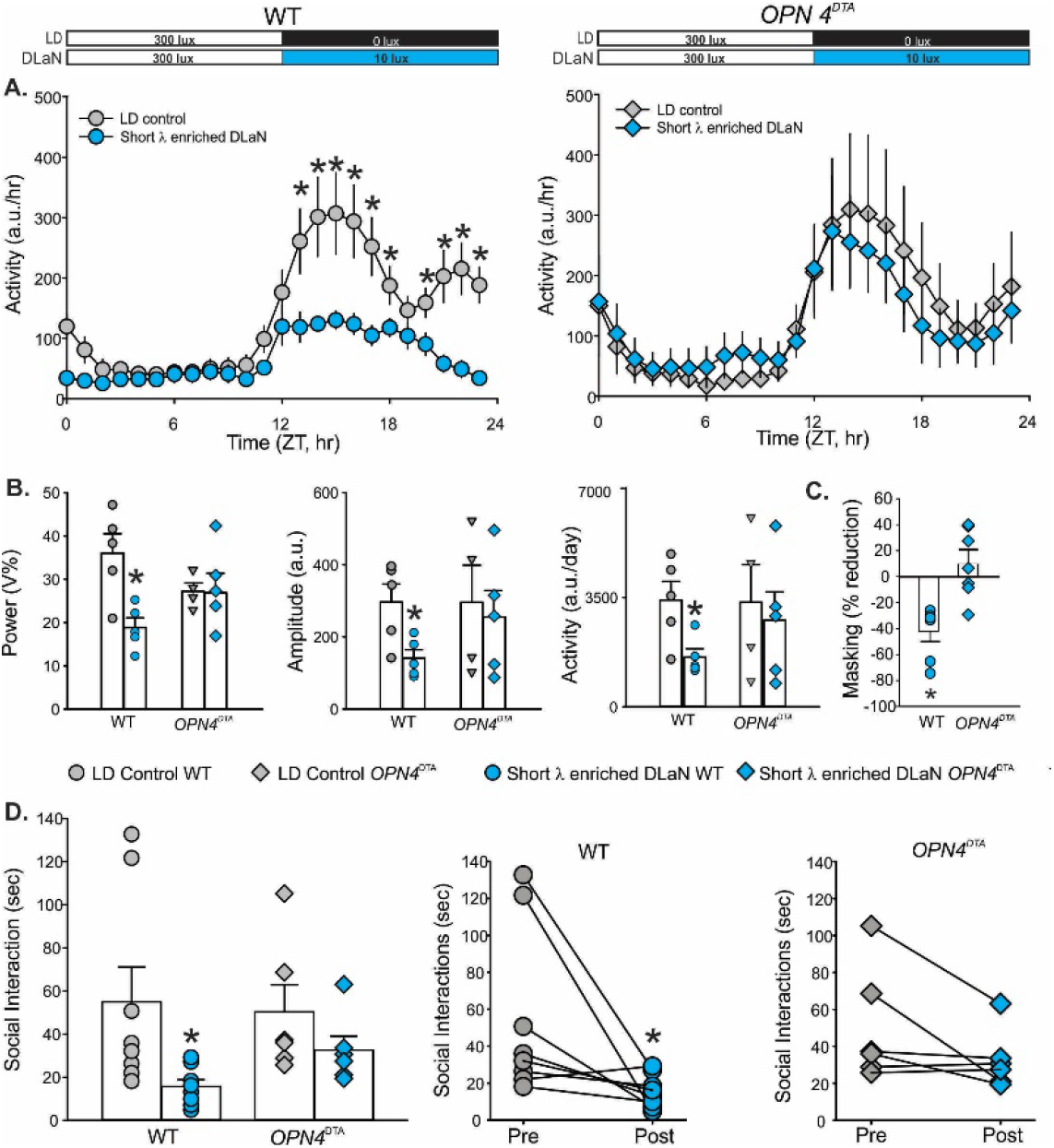
Melanopsin mediates the impacts of DLaN on activity rhythms and social behavior. WT and *Opn4*^DTA^ were kept in baseline conditions (LD) or exposed to the short λ enriched DLaN (10 lx) for two weeks. (**A**) Waveforms of daily rhythms in cage activity of WT (left) or *Opn4*^DTA^ mice (right) under LD control conditions or under the short λ enriched DLaN. The activity waveform (1 hr bins) of each group was analyzed using a two-way ANOVA with treatment and time as factors. The WT mice exhibited a significant effect of treatment (F_(1, 239)_ = 95.39; *P* < 0.001), time (F_(23, 239)_ = 11.674; *P* < 0.001) as well as a significant interaction between the two factors (F_(23, 239)_ = 3.047, *P* < 0.001). In contrast, the *Opn4*^DTA^ mice exhibited a significant effect of time (F_(23,215)_ = 4.73; *P* < 0.001) but not treatment (F_(1, 215)_ = 0.636, *P* = 0.426). There was not any evidence for an interaction (F_(23,215)_ = 0.611; *P* = 0.917). Asterisks indicate significant (*P* < 0.05) differences between the 1 hr bins as measured by Holm-Sidak test for multiple comparisons. **(B)** Analyses of the properties of the daily activity rhythms (10 day recordings) including power, amplitude, and overall activity levels. **(C)** Light-evoked negative masking was measured by exposing the mice to the short λ enhanced light (10 lx) for 1h at ZT14 and comparing their activity to that of the prior day. **(D)** The impact of the short λ enhanced DLaN on WT and *Opn4*^DTA^ social interactions was measured by comparing the behavior of the mice in baseline conditions (LD) and then after 2 weeks exposure to the short λ enhanced DLaN. A paired *t*-test was used to analyze the changes before and after light exposure (**Table 2**) with the asterisks indicating significant difference (*P* < 0.05). Histograms show the means ± SEM with the values from individual animals overlaid. a.u.= arbitrary units

### Reduced melanopic illumination prevents DLaN-driven disruption of daily rhythms in activity in WT and Cntnap2 KO mice

Next, we used the LED array to determine whether long λ enriched illumination would reduce the DLaN disruption of locomotor activity rhythms. Cohorts of *Cntnap2* KO and WT mice were first housed under control conditions (LD 12:12), then randomly exposed to either long or short λ enriched DLaN (10 lx) for 2 weeks and the waveforms examined (**Fig. 3A)**. Under baseline conditions, both *Cntnap2* KO mice and WT controls exhibited robust rhythms in activity, although several parameters (total activity, power, amplitude) measured from the *Cntnap2* KO mice were consistent with weaker rhythms as previously described (Wang et al., 2020). When treated with the short λ enriched DLaN, the average waveform (1 hr bins) of both genotypes was significantly altered (**Fig. 3A, B; Table 3)** The WT and the *Cntnap2* KO mice exhibited a significant reduction in a number of activity rhythm parameters while the long λ enriched illumination did not produce significant changes compared to baseline (**Fig. 3; Table 3)**. These differences in the effects of the short and long λ illumination were strikingly evident when some of the key activity parameters were examined before and after treatment (**Fig. 4**). These results indicate that the negative effects of DLaN are driven by the spectral properties of the light used for illumination in both WT and *Cntnap2* KO mice.

**Fig. 3:**
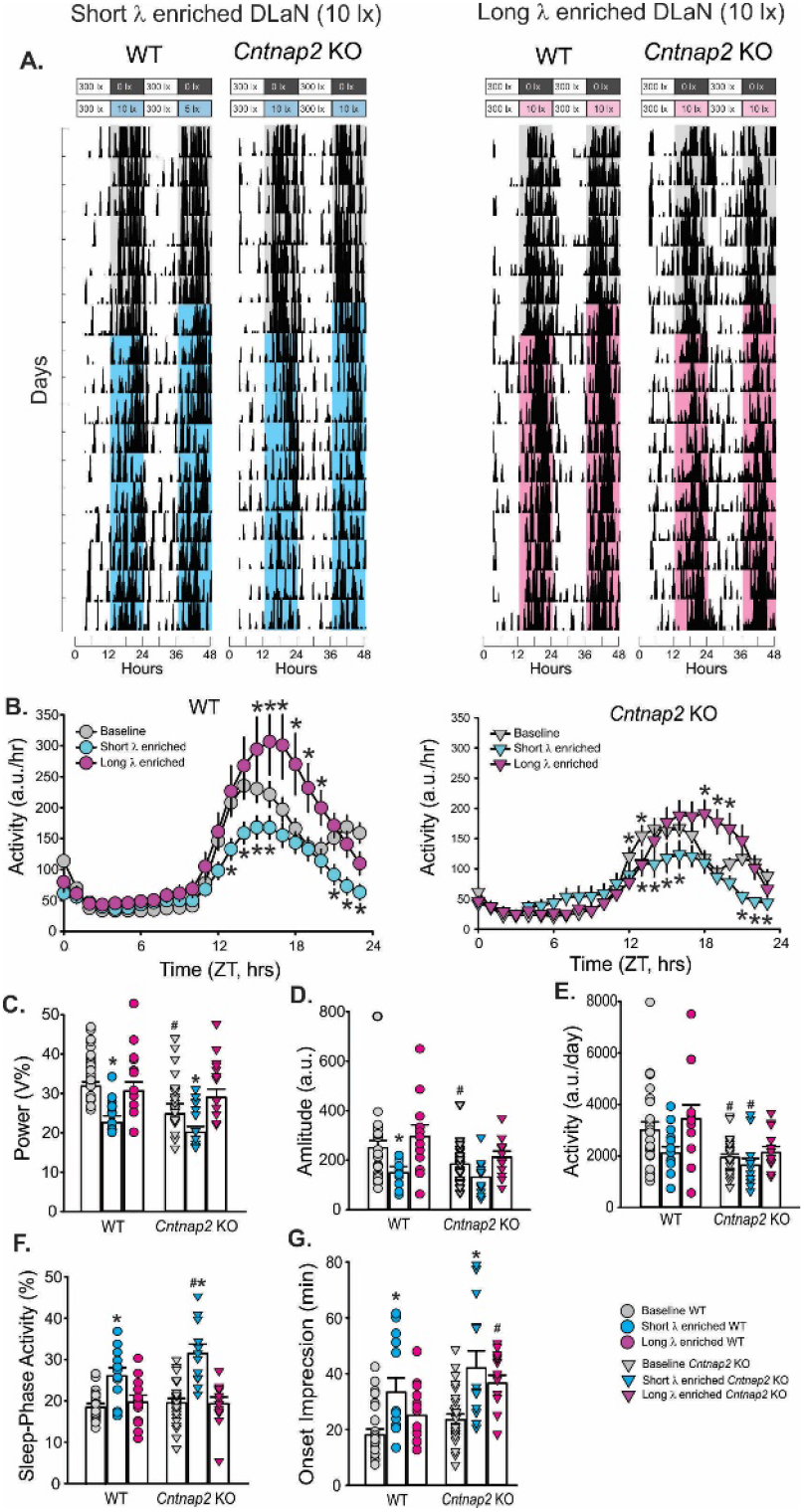
Long wavelength illumination minimizes DLaN disruption of daily activity rhythms in WT and *Cntnap2* KO mice. **(A)** Examples of actograms of daily rhythms in cage activity of WT and *Cntnap2* KO mice exposed to a short λ (left) or a long λ (right) enriched DLaN (10 lx) for 2 weeks. The activity levels in the actograms were normalized to the same scale (85% of the maximum of the most active individual. Each row represents two consecutive days, and the second day is repeated at the beginning of the next row. The bars on the top of actograms indicates the time and intensity when the nightly lights were given. The cyan shading represents the short λ enhanced lighting and the magenta shading the long λ enhanced lighting. **(B)** Waveforms of daily rhythms in cage activity under short (teal) or long (magenta) λ enriched DLaN in WT (circles) and *Cntnap2* KO (triangles) mice. The baseline values from mice under control conditions (LD) are shown for comparison (grey). The activity waveform (1 hr bins) of each group was analyzed using a Two-way ANOVA for repeated measures with treatment and time as factors. Both groups exhibited significant effects of time (WT: F_(23,1151)_ = 43.29, *P* < 0.001; *Cntnap2* KO: F_(23,1151)_ = 44.96, *P* < 0.001) and treatment (WT: F_(2,1151)_ = 42.66, *P* < 0.001; *Cntnap2* KO: F_(2,1151)_ = 15.16, *P* < 0.001), and the interaction between the two factors was identified (WT: F_(46,1151)_ = 2.57, *P* < 0.001; *Cntnap2* KO: F_(46,1151)_ = 3.8, *P* < 0.001). Asterisks indicate significant (*P* < 0.05) differences between the 1 hr bins as measured by Holm-Sidak test for multiple comparisons. **(C-G)** Properties of the daily activity rhythms (10 day recordings) in baseline conditions (LD) and under short or long λ enhanced DLaN were analyzed using Two-way ANOVA with genotype and treatment as factors followed by the Holm-Sidak test for multiple comparisons (n=12 per genotype/treatment group; see also **Table 3**). Histograms show the means ± SEM with the values from the individual animals overlaid. Significant (*P* < 0.05) differences as a result of treatment or genotype are indicated with an asterisk or a crosshatch, respectively. a.u.= arbitrary units

**Fig. 4:**
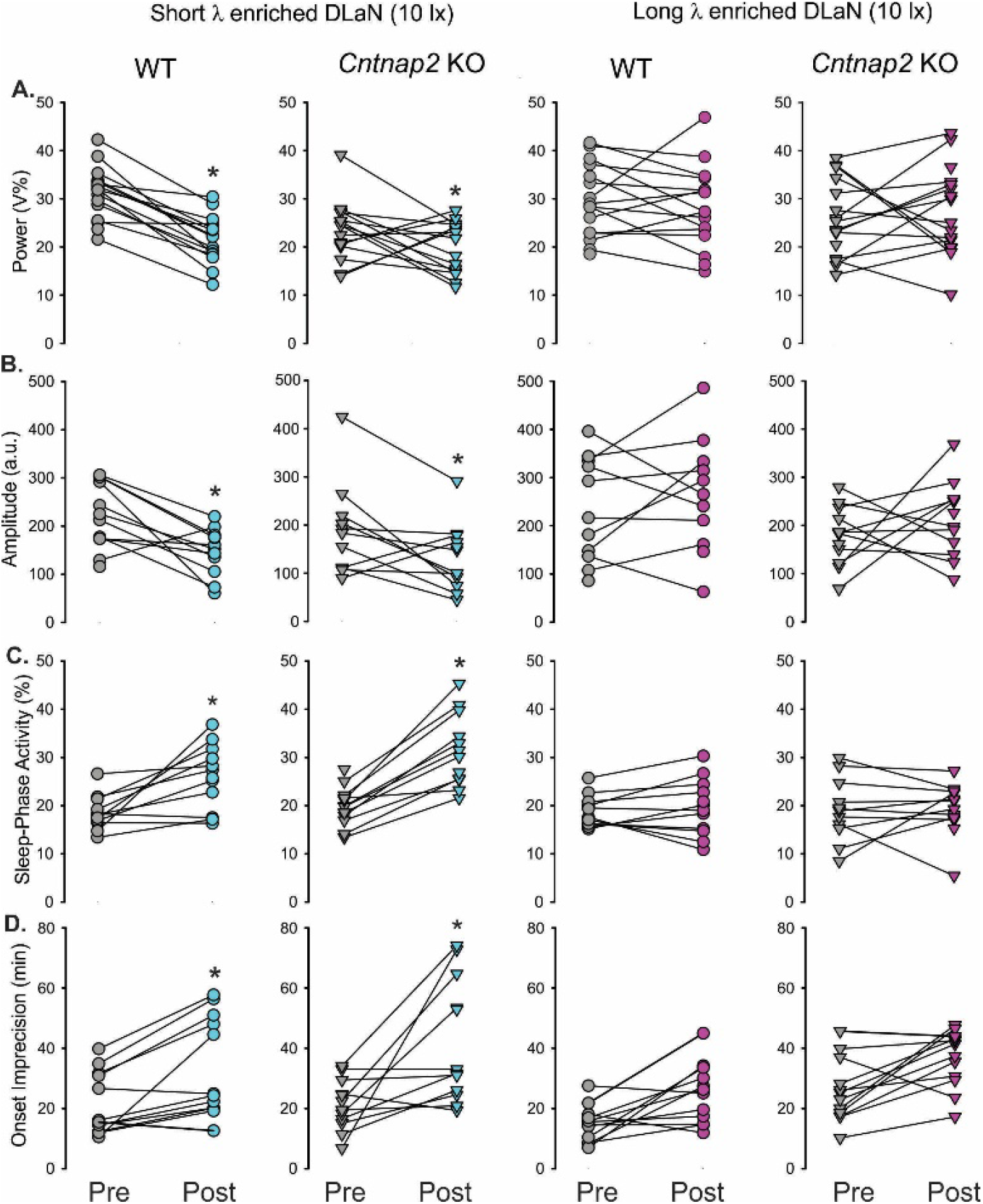
Long wavelength illumination was less disruptive to activity rhythms in WT and *Cntnap2* KO mice. Key parameters of the daily activity rhythms were measured before (grey) and after exposure to the short λ (teal) or the long λ (magenta) enriched DLaN treatment. Values from individual WT (circles) and *Cntnap2* KO (triangles) mice are reported. A paired *t*-test was used to analyze the changes before and after the treatment, and an asterisk over the treated values indicates significant difference (*P* < 0.05).

### Reduced melanopic illumination prevents DLaN-driven disruption of social behavior in WT and Cntnap2 KO mice

DLaN reduced social behavior in both *Cntnap2* KO mutants as well as WT controls (Wang et al., 2020), thus, here we sought to determine the influence of the lighting treatments using a reciprocal social interaction test. The short λ enriched DLaN treatment decreased social behavior in both *Cntnap2* KO and WT mice, while the long λ enriched light was without effect (**Fig. 5, Table 3**). The short λ DLaN decreased social interactions (>10%) in 10 out of the 12 mutant mice evaluated, whereas the long λ reduced social interactions in 5 out of 12 KO mice (**Fig. 5B, C**). Our findings indicate that exposure to short λ enriched DLaN disrupted social behavior in both genotypes, and the use of longer λ illumination could effectively reduce the impact of DLaN on social behavior.

**Fig. 5:**
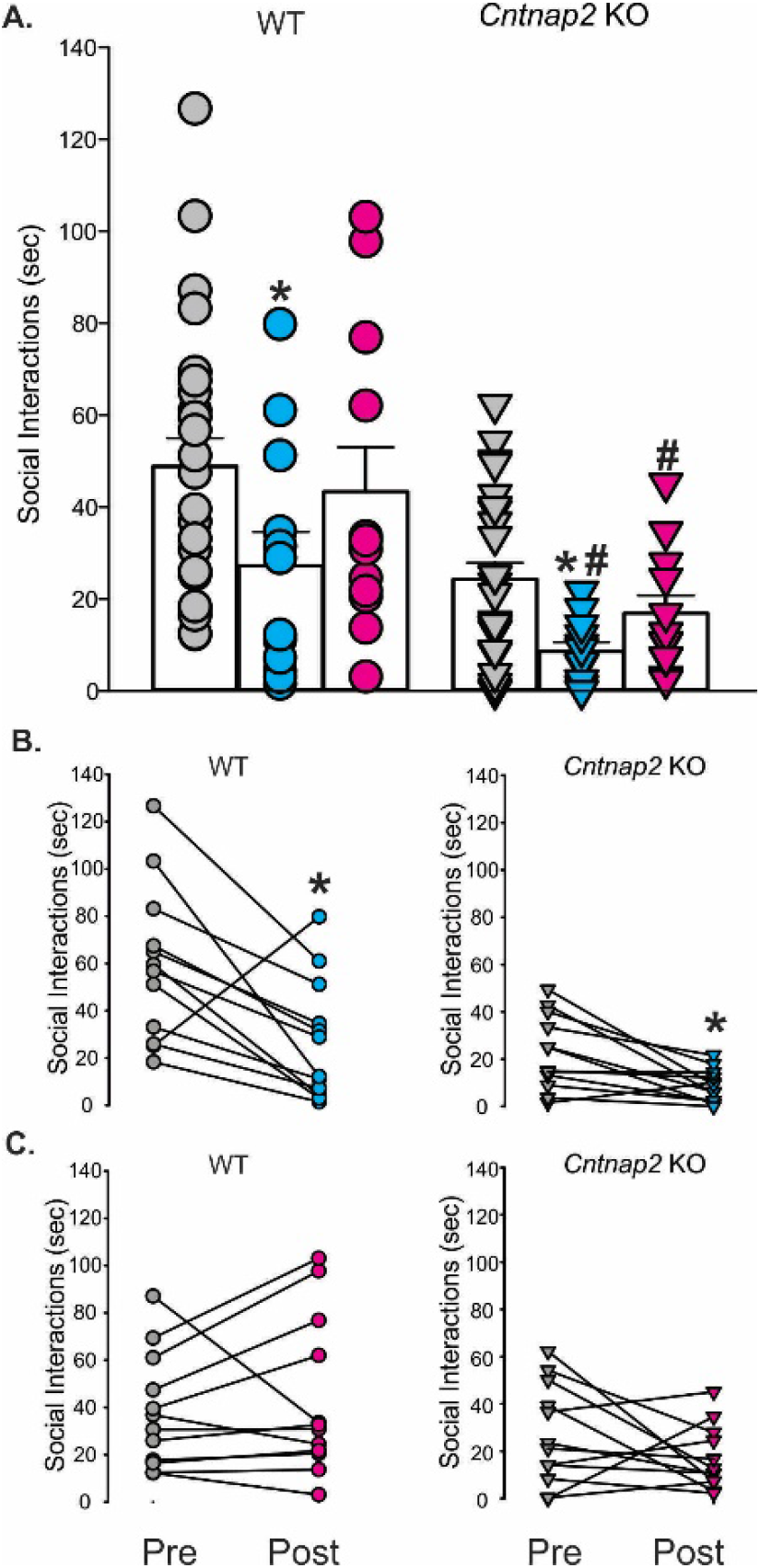
Long wavelength illumination diminished DLaN-evoked social deficits in WT and *Cntnap2* KO mice. The social behavior was evaluated by analyzing the time actively spent by the testing mouse in social interactions with the novel stranger mouse. The stranger and testing mice were matched for age, sex and genotype. Experiments were conducted during the active phase at ZT 18. **(A)** Time spent in social interactions. Histograms show the means ± SEM with overlaid the values from individual WT (circles) and *Cntnap2* KO (triangles) mice in LD (grey) and under the short λ (teal) or the long λ (magenta) enriched DLaN. Data were analyzed using a two-way ANOVA with genotype and treatment (see **Table 3**), followed by the Holm-Sidak multiple comparisons test. The asterisks indicate significant differences (*P* < 0.05) between LD and DLaN treatments within genotype and the crosshatches between genotypes. **(B, C)** Values from individual WT (circles) and *Cntnap2* KO (triangles) animals before (grey) and after exposure to the short λ (teal) or long λ (magenta) enriched DLaN. Data were analysed using a paired *t*-test. The asterisks indicate significant difference (*P* < 0.05).

### Reduced melanopic illumination prevents DLaN-driven disruption of repetitive behavior in Cntnap2 KO mice

We have previously shown that DLaN increases repetitive behavior in *Cntnap2* KO mutants but not in WT (Wang et al., 2020), hence, we explored the effects of the two lighting treatments on grooming behavior. As previously observed, the *Cntnap2* KO exhibited higher levels of grooming than WT mice under baseline conditions, and neither DLaN treatment altered the levels of grooming in WT mice (**Fig. 6, Table 3)**. The short λ enriched DLaN treatment triggered a selective increase in the time of grooming in the *Cntnap2* KO mice (**Fig. 6, Table 3**). In particular, the short λ DLaN increased grooming (>10%) in 16 out of the 22 KO mice evaluated while the long λ DLaN only affected 4 mutants (**Fig. 6B, C**). Our observations demonstrate that the impact of DLaN on repetitive grooming in the *Cntnap2* KO was dependent upon the λ used and that the mutants are selectively vulnerable to the effects of DLaN on repetitive behavior.

**Fig. 6:**
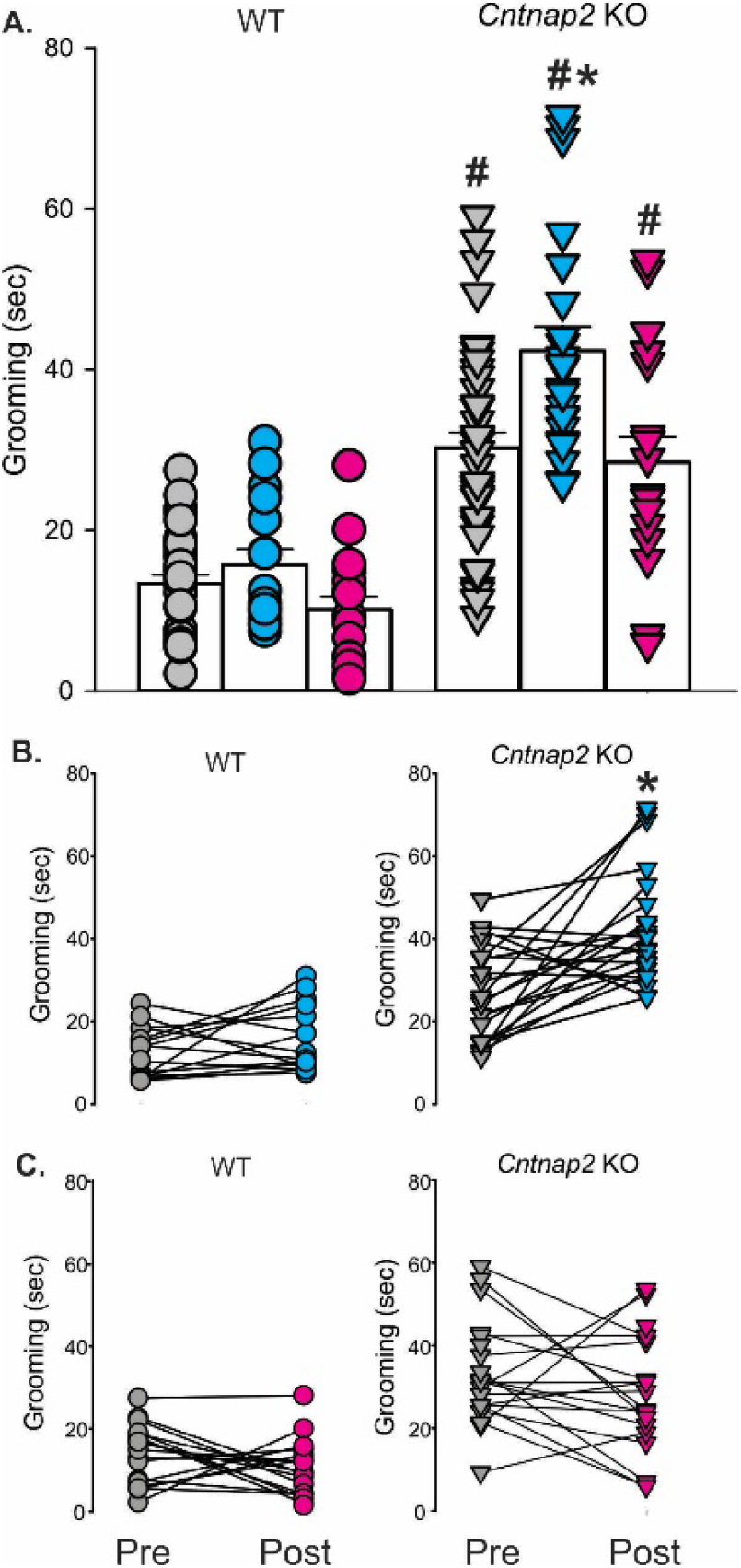
Long wavelength illumination DLaN lessened repetitive behavior in *Cntnap2* KO mice. Grooming was assessed in a novel arena in mice of each genotype under each lighting condition (baseline LD, short λ and long λ enriched DLaN). Measurements were conducted at ZT 18. **(A)** Time spent grooming by WT (circles) and *Cntnap2* KO (triangles) mice under LD (grey), short λ (teal) or long λ (magenta) enriched DLaN. Histograms show the means ± SEM with the values from the individual animals overlaid. Data were analyzed using a Two-way ANOVA with genotype and treatment as factors (see **Table 3**) followed by the Holm-Sidak multiple comparisons test. Significant (*P* < 0.05) effects of treatment or genotype are indicated with an asterisk or a crosshatch, respectively. **(B, C)** Values from individual WT and *Cntnap2* KO mice animals before (grey) and after exposure to the short λ (teal) or long λ (magenta) enriched DLaN. A paired *t*-test was used to analyze the changes before and after the treatment, and an asterisk over the treated values indicates significant difference (*P* < 0.05).

### Limited Sex differences in the DLaN-driven behavioral changes in WT and Cntnap2 KO mice

Given the strong male bias in prevalence of ASD, we powered our behavioral studies to detect possible sex difference in behavior. First, we compared key measures of activity rhythms, social interactions and grooming between the two genotypes in mice held in the LD cycle. Most activity parameters varied with genotype but not sex (**Table 4**) with the *Cntnap2* KO showing hypo-activity throughout the night. As has been described previously, the activity waveforms were more robust in the WT females compared with males with higher activity levels in the early night (**Supplemental Fig. 1**). We found a similar situation for social behavior where there were significant effects of genotype but not sex (**Table 4**). By contrast, for grooming behavior there were significant effects of both genotype and sex with the female mutants showing significantly altered measures of repetitive behavior (**Table 4**). Next, we compared these same parameters in male and female WT and *Cntnap2* KO mice held under short λ enriched DLaN conditions. Under these conditions, there were no significant sex differences in any of the behaviors. For activity measures, there were not even differences between the genotypes as all of rhythms were compromised (**Table 5; Supplemental Fig. 2)**. For social interactions and grooming, both sexes were compromised more in the mutant mice (**Table 5**). In short, although we powered our sample size to detect possible sex differences, for the most part, both sexes were impacted by the *Cntnap2* mutation. As measured by grooming, the repetitive behavior is not as robust a phenotype in the female mutants.

**Table 5:** Sex differences in WT and *Cntnap2* KO mice under DLaN conditions. Comparisons of age-matched male and female WT and *Cntnap2* KO mice held under short wavelength DLaN conditions for two weeks. Values are shown as averages ± SEM. Data were analyzed with a two-way ANOVA using genotype and treatment as factors. There were no significant interactions between genotype and sex, thus, these are not reported. The Holm-Sidak test for multiple comparisons was used when appropriate. The hashtags indicate significant differences between genotypes. Significant differences are shown in bold. a.u.= arbitrary units

| Measures | WT |  | *Cntnap2* KO |  | Two-way ANOVA |  |
| --- | --- | --- | --- | --- | --- | --- |
|  | Male DLaN<br>n=6 | Female DLaN<br>n=6 | Male DLaN<br>n=6 | Female DLaN<br>n=6 | Genotype | Sex |
| Total activity (a.u./24 hrs) | 1755.3 ± 257.6 | 2473.4 ± 359.5 | 2152.9 ± 388.3 | 1892.6 ± 216.5 | $F_{(1,23)}=0.08$; $P=0.773$ | $F_{(1,23)}=0.53$; $P=0.474$ |
| Power (% variance) | 21.0 ± 1.0 | 24.2 ± 1.6 | 25.2 ± 1.8 | 22.0 ± 0.9 | $F_{(1,23)}=0.52$; $P=0.479$ | $F_{(1,23)}=0.01$; $P=0.999$ |
| Amplitude (Peak/trough) | 135.7 ± 19.5 | 161.7 ± 20.4 | 187.3 ± 25.7 | 156.8 ± 20.0 | $F_{(1,23)}=1.18$; $P=0.291$ | $F_{(1,23)}=0.01$; $P=0.919$ |
| Social interaction (sec) | 38.1 ± 11.3 | 17.8 ± 7.7 | **7.9 ± 3.0<sup>#</sup>** | **11.0 ± 2.1<sup>#</sup>** | $F_{(1,23)}=6.20$; $P=0.022$ | $F_{(1,23)}=1.35$; $P=0.259$ |
| Grooming (sec) | 18.0 ± 2.7 | 13.42 ± 2.8 | **44.4 ± 3.2<sup>#</sup>** | **44.9 ± 4.5<sup>#</sup>** | $F_{(1,37)}=62.23$; $P<0.001$ | $F_{(1,37)}=0.31$; $P=0.578$ |

### Reduced melanopic illumination diminished DLaN induction of c-Fos in the basolateral amygdala in Cntnap2 KO mice

Prior work has shown that activation of the ipRGC can drive cFos expression in several brain regions including the hypothalamus and the limbic system (Milosavljevic et al., 2016; Fernandez et al., 2018). Hence, to unravel some of the possible mechanisms underlying the impacts of DLaN on autistic behaviors, we examined c-Fos immunoreactivity across different brain regions known to be implicated in the control of repetitive behavior, such as the BLA, Periventricular Nucleus (PVN), and Lateral Hypothalamus (LH). For controls, we also examined possible DLaN driven c-Fos expression to two areas known to receive light information from ipRGC including the SCN and peri-Habenular nucleus (pHb). Compared to mice held on a standard LD cycle, WT and *Cntnap2* KO mice exposed for 2 weeks to the short λ enriched DLaN displayed a significantly increased number in c-Fos positive cells in both the BLA and pHb, but not in other brain regions of interest (**Fig. 7, Table 6**). In contrast, the long λ enriched DLaN did not induced c-Fos expression in any of these regions. Prior work suggested that the glutamatergic cell population in the BLA is a critical part of a circuit responsible for repetitive behavior (Felix-Ortiz et al., 2016). Therefore, we crossed the *Cntnap2* KO line into mice that expressed both vGAT2-Cre (Vong et al., 2011) and a Cre-dependent reporter ZsGreen1 (Ai6) (Madisen et al., 2010), which provides strong cell body labelling. WT and *Cntnap2* KO mice with these reporters were then exposed to short λ enriched DLaN for two weeks. The expression of the vGlut2-ZsGreen1 reporter within the BLA region was robust in both the WT and mutants. Most importantly, the short λ DLaN elicited a higher number of vGlut2-ZsGreen1/cFos immunopositive in the *Cntnap2* KO mice (**Table 7**) in comparison to the WT under DLaN or the WT and mutants under LD conditions. Both genotypes exhibited a significantly higher co-expression in the light treated group (**Table 7**). These data suggest that the glutamatergic cells within the BLA are activated by DLaN and raise the possibility that this cell population contributes to its effect on repetitive behavior.

**Table 6.** Long wavelength enriched illumination minimized the DLaN evoked c-Fos expression in several brain regions. Number of c-Fos immunopositive cells in age-matched WT and *Cntnap2* KO mice under LD or DLaN regimen (n = 6/group). Immunopositive cells were counted in the left and or right region of interest in two consecutive slice and the numbers averaged to obtain one value per animal. Values are shown as averages ± SEM. Data were analyzed with a two-way ANOVA using genotype and treatment as factors. The Holm-Sidak test for multiple comparisons was used when appropriate. There were no significant interactions between genotype and treatment. The asterisks indicate significant differences between LD and DLaN treatments within genotype. SCN=Suprachiasmatic Nucleus; pHb=perihabenular nucleus; PVN=Periventricular Nucleus; LH=Lateral Hypothalamus; BLA=BasolateralAmygdala. Significant differences are shown in bold.

| Brain region | WT |  |  | *Cntnap2* KO |  |  | Two-Way ANOVA |  |
| --- | --- | --- | --- | --- | --- | --- | --- | --- |
| | Baseline LD | Short $\lambda$ DLaN | Long $\lambda$ DLaN | Baseline LD | Short $\lambda$ DLaN | Long $\lambda$ DLaN | Genotype | Treatment |
| SCN | 4.6 ± 1.3 | 4.1 ± 0.8 | 4.6 ± 0.9 | 4.8 ± 1.5 | 3.3 ± 0.7 | 4.5 ± 0.7 | $F_{(1,35)}=0.09; P=0.77$ | $F_{(2,35)}=0.62; P=0.54$ |
| pHb | 30.1 ± 7.1 | **55.4 ± 6.7\*** | 34.7 ± 4.9 | 22.2 ± 1.6 | **63.1 ± 8.1\*** | 38.0 ± 6.4 | $F_{(1,35)}=0.047; P=0.83$ | **$F_{(2,35)}=71.94; P<0.001$** |
| PVN | 28.4 ± 4.4 | 35.5 ± 4.8 | 29.6 ± 6.9 | 35.8 ± 7.6 | 35.5 ± 3.9 | 28.0 ± 7.0 | $F_{(1,35)}=0.19; P=0.67$ | $F_{(2,35)}=0.76; P=0.48$ |
| LH | 134.1 ± 42.0 | 133.6 ± 30.2 | 150.9 ± 28.2 | 140.1 ± 40.0 | 140.9 ± 12.5 | 121.7 ± 19.9 | $F_{(1,35)}=0.055; P=0.82$ | $F_{(2,35)}<0.001; P=1.00$ |
| BLA | 170.2 ± 38.1 | 235.1 ± 46.3 | 160.2 ± 29.9 | 202.0 ± 42.0 | **346.3 ± 27.1\*** | 222.9 ± 25.8 | **$F_{(1,35)}=7.99; P=0.008$** | **$F_{(2,35)}=5.99; P=0.006$** |

**Table 7.** Increased c-Fos expression in glutamatergic neurons in the BLA by the short wavelength enriched illumination. Comparisons of vGlut2-ZsGreen1/c-Fos positive cells in the BLA of WT and *Cntnap2* KO mice under LD or DLaN regimen (n = 4/group) showed a marked increase in the percentage of c-Fos positive in the mutants exposed to short wavelength enriched DLaN. The number of vGlut2-ZsGreen1 expressing neurons and, among these neurons, the number of c-Fos positive was determined in 4 to 7 images per animal. The percentage of co-expressing cells was then calculated per image. These values were averaged to obtain a single number per animal. Data are shown as the average ± SEM and were analyzed with a two-way ANOVA using genotype and treatment as factors followed by the Holm-Sidak test for multiple comparisons. The asterisks indicate significant differences between LD and DLaN treatments within genotype. Significant differences are shown in bold.

|  | vGlut2-ZsGreen1 cells | vGlut2-ZsGreen1/c-Fos positive | % co-expression |
| --- | --- | --- | --- |
| WT LD | 43.4 ± 3.1 | 9.1 ± 0.7 | 21.0 ± 1.0 |
| WT + DLaN | 47.2 ± 2.2 | 18.9 ± 1.4 | **40.5 ± 4.0\*** |
| *Cntnap2* KO LD | 45.2 ± 2.3 | 8.8 ± 1.8 | 19.7 ± 4.6 |
| *Cntnap2* KO + DLaN | 33.2 ± 1.0 | 17.7 ± 1.3 | **53.6 ± 5.5\*** |
| **Two-Way ANOVA** | Treatment | Genotype | Interaction |
| | $F_{(1, 12)}=41.68$; $P<0.0001$ | $F_{(1,12)}=2.017$; $P=0.181$ | $F_{(1, 12)}=2.998$; $P=0.109$ |

**Fig. 7:**
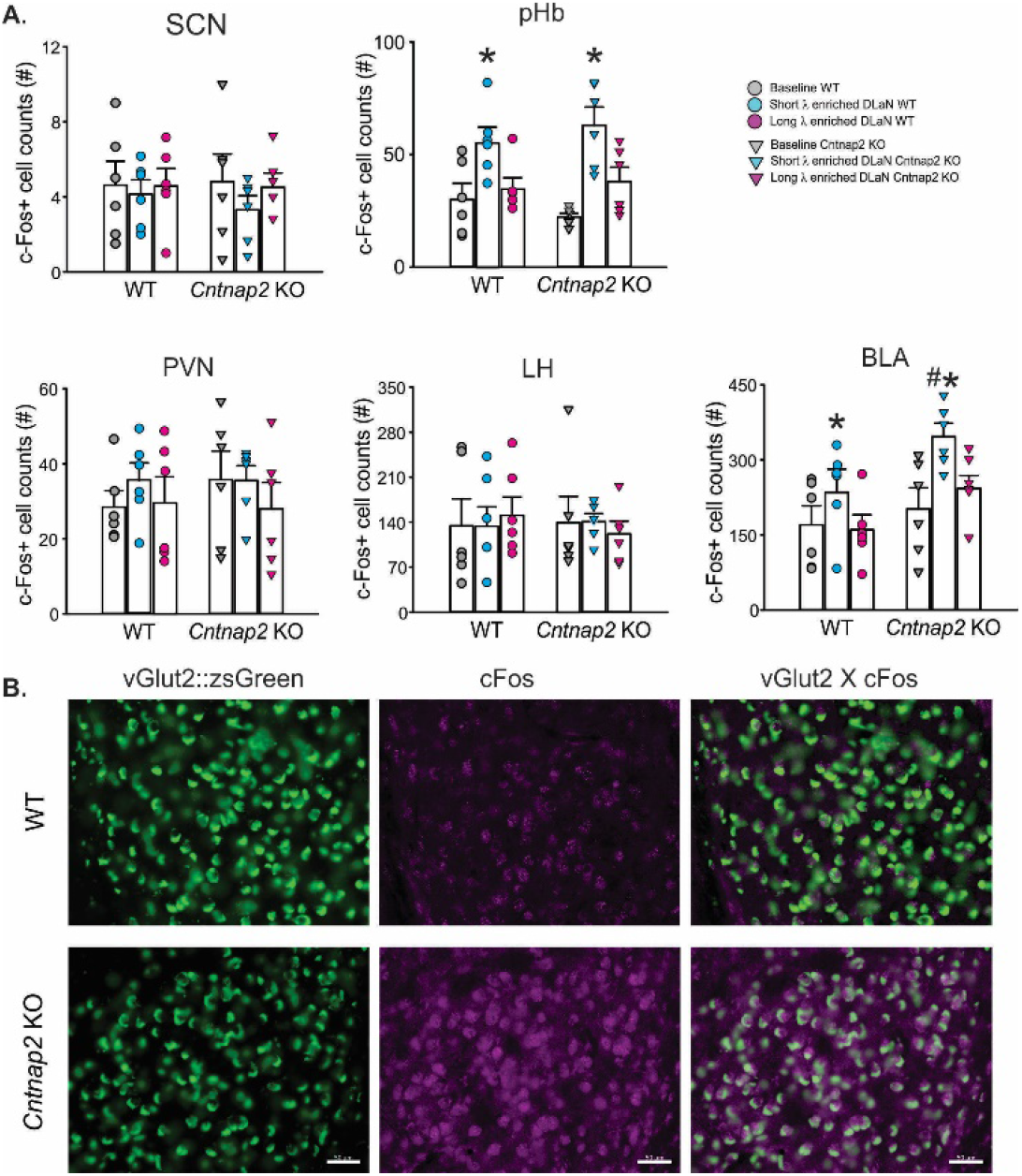
The short, but not the long, wavelength illumination elicits a selective increase in c-Fos immunopositive cells in the basolateral amygdala. Brains were collected at ZT 18 from WT (circles) and *Cntnap2* KO (triangles) mice held in LD or exposed to the short λ (teal) or the long λ (magenta) enriched DLaN (10 lx) for two weeks. (**A**) The number of c-Fos positive cells was determined bilaterally in several brain regions including the SCN, the peri-Habenula (pHb), the paraventricular nucleus (PVN), the lateral hypothalamus (LH) and the basolateral amygdala (BLA). The values from each region of interest (both left and right hemisphere, two consecutive slices/animal) were averaged to obtain one value per animal (n=6), and are shown as the mean ± SEM (histograms with the values from the individual animals overlaid). Data were analyzed with a two-way ANOVA with genotype and treatment as factors (see **Table 6**) followed by Holm-Sidak multiple comparison test. Significant (*P* < 0.05) effects of treatment or genotype are indicated with an asterisk or a crosshatch, respectively. (**B**) Exposure to the short λ enriched DLaN for two weeks activated more glutamatergic neurons in the mutants in comparison to the WT. Representative images of vGlut2-ZsGreen1 neurons immunopositive for c-Fos in the BLA of WT and *Cntnap2* KO mice heterozygous for vGlut2-ZsGreen1. WT and *Cntnap2* KO were perfused at ZT18 and the brains collected. The total number of vGlut2-ZsGreen1 neurons and, among these, of those immunopositive for c-Fos was determined in 4 to 7 images acquired at random from 6 consecutive slices per animal (n=4). The values from each image per animal were averaged and analyzed with a two-way ANOVA followed by the Holm-Sidak multiple comparisons test (see **Table 7**). Scale bars=50 μm.

## Discussion

Nightly light exposure is a common environmental perturbation and has been shown to cause a number of ill effects (Lucassen et al., 2016; Bumgarner et al., 2021). Some populations may be more vulnerable to the negative effects of the exposure of light at night. For example, in the *Cntnap2* KO mouse model of ASD, we have found that DLaN selectively increases repetitive behavior as measured by grooming while WT mice were not impacted (Wang et al., 2020). On the other hand, DLaN negatively impacted other behaviors including social interactions and daily activity rhythms in both the *Cntnap2* KO and WT mice. In the present study, we confirmed these negative effects of nocturnal lighting on activity rhythms, social interactions, and repetitive behaviors (**Table 3**). Logically, the simplest approach to reducing these negative effects would solely be to decrease light intensity at night while ensuring a robust exposure to sunlight during the day. However, according to surveys, people spend a majority of time indoors (as high as 90%) and have need for illumination after the sun sets (e.g. Klepeis et al., 2001). Therefore an appealing alternative approach would be to adjust the spectral quality of indoor lighting to minimize the negative effects. As a test case, in the present study, we used an array of LEDs to generate dim illumination (10 lx) to determine whether the spectral composition of nightly illumination would make a difference in the *Cntnap2* KO model of autism.

The retinal photoreceptor system has unique intensity and wavelength dependent characteristics (Lucas et al., 2014). The negative effects of nightly light exposure are likely to be mediated by ipRGCs expressing the photopigment melanopsin, which is maximally sensitive to light with a peak response to ≈480nm light (Hattar et al., 2002; Panda et al., 2005). Indeed a number of studies have found evidence that short λ enriched lighting is more effective for circadian responses (for review see Brown, 2020). However, recent work emphasizes that ipRGCs also receive input from rod and cone photoreceptors (Hattar et al., 2003; Lucas et al., 2012; Van Diepen et al., 2021; Schoonderwoerd et al., 2022) with rods driving the circadian response to low-intensity light (Altimus et al., 2010; Lall et al 2010) like the 10 lx used in the present study. Here we used an LED lighting system (Korrus Inc.) to tailor the spectral properties of the DLaN with the goal of manipulating melanopic stimulation. The output of the LEDs was measured as power over a defined wavelength (380-780nm) using a spectrophotometer. This data was analysed using a CIE toolkit which enabled us to estimate that the short λ illumination was at 15 melanopic lx while the long λ illumination was 1 melanopic lx while maintaining illuminance at ≈10 lx.

Using these two lighting treatments with predicted high and low melanopic stimulation, we measured the impact of the illumination on three classic tests of the ipRGC input to the circadian system (**Fig. 1**). We demonstrated that the low melanopic stimulation reduced the effect of light on negative masking, light-induced phase shifts and light-induction of c-Fos in the SCN. Next, we examined the influence of the high melanopic illumination on activity and social behavior of WT and *Opn4*^DTA^ in which the ipRGCs are lost due to targeted diphtheria toxin expression. We found that the high melanopic illumination showed greatly diminished ability to disrupt activity rhythms including negative masking as well as alter social behavior in the mutant mice (**Fig. 2**). Together, this data indicates that ipRGCs are required for a majority of the negative effects of DLaN and that controlling melanopic stimulation altered activation of the ipRGC/SCN pathway.

With this validated lighting system in hand, we sought to use the LED array to determine whether long λ enriched illumination would diminish DLaN disruption of behavior in WT and *Cntnap2* KO mice. Both lines of mice exhibited a significant reduction in a number of activity rhythm parameters when exposed to the high melanopic illumination while the low melanopic illumination did not produce significant changes compared to baseline (**Fig. 3, 4**). Similarly, exposure to high melanopic illumination disrupted social behavior in both genotypes while the low melanopic DLaN had little effect on social interactions (**Fig. 5**). Finally, we found that exposure to high melanopic illumination increased repetitive grooming in the *Cntnap2* KO but not the low melanopic DLaN (**Fig. 6**). Together, these data show that minimizing the melanopic activation of the light source was an effective strategy to minimize the negative effects of night-time light exposure.

There is a strong male bias in ASD prevalence with a common reported ratio of ∼4:1 (Fombonne, 2009) and there is evidence for significant interactions between sex and genotype in the *Cntnap2* gene in human populations (e.g. Hsu et al., 2019). Some of these sex differences may be driven in part by male/female differences in phenotypic presentation, including fewer repetitive behaviors in females (e.g. Werling and Geschwind, 2013). On the other hand, sleep disturbances appear to be more common in female with ASD (Elkhatib Smidt et al., 2021). Prior work with the *Cntnap2* KO mouse model found clear evidence for a sex difference with home cage activity with robust hypo-activity in the males but not in the female mutants (Angelakos et al., 2019). In addition, the functional responses of cortical circuitry in male mice are more strongly affected by *Cntnap2* mutations than females (Townsend and Smith, 2017). Therefore, we powered the activity analysis to be able to examine possible sex differences in activity rhythms. Sex differences could be observed in the activity rhythms in WT mice that were lost in the *Cntnap2* KO line (**Table 4; S. Fig. 1**). We did see evidence that the excessive grooming, a measure of repetitive behavior, was more pronounced in the mutant males than in the females, although both sexes showed significantly higher grooming than age and sex-matched WT. Social deficits were observed in mutants of both sexes. Hence, both sexes of *Cntnap2* KO exhibited phenotypes of disrupted activity rhythms, higher repetitive behavior and less social interactions compared to WT mice. Interestingly, prior work has found that LPS-induced maternal immune activation caused male-specific deficits in behavior and gene expression in the *Cntnap2* KO line (Schaafsma et al., 2017). Furthermore, we asked whether the female mice might be less influenced by the environmental challenge caused by DLaN treatment. In our three assays (activity, grooming, and social interactions), we found that both sexes were negatively impacted by the DLaN (**Table 5; S. Fig. 2**). The females *Cntnap2* KO did appear to be relatively resistant to the excessive grooming triggered by DLaN. Still overall our data demonstrate that the both sexes were negatively impacted by the Cntnap2 mutation and the negative influence of DLaN.

Prior work has shown that activation of ipRGC can drive c-Fos expression in several brain regions including those in the hypothalamus and limbic system (Milosavljevic et al., 2016; Fernandez et al., 2018). We sought to determine whether the 2-weeks of DLaN would alter c-Fos expression in some of those brain regions (**Fig. 7**). In contrast to the effects of the light when the mice were held in DD (**Fig. 1**), we did not see any evidence that the chronic nightly high melanopic illumination increased c-Fos in the SCN. Similarly, we did not see increased c-Fos expression in the PVH and LH. Interestingly, the chemogenetic activation of ipRGCs, did induced c-Fos expression in these two regions (Milosavljevic et al., 2016). The difference may well be the relatively weak light stimulus used in the present study or the difference between acute chemogenetic activation vs. chronic stimulation with the DLaN. We did find that our high melanopic DLaN did increases c-Fos expression in the pHb and BLA. Prior work has implicated the pHb as a region critically involved in the effects of light on mood (Fernandez et al., 2018).

Specifically, there is support for circuit by which the ipRGC regulation of the dorsal region of the pHb projects to the nucleus accumbens (NAc) to drive depressive-like behaviors (An et al., 2020). Thus, we would expect to see DLaN driven alterations in depressive behaviors in our mouse models (Fonken et al., 2012; Bedrosian et al., 2013; Walker et al., 2020) and this will be an important area for future work. More critical for the present study, prior work suggests that the BLA may regulate repetitive behavior (Felix-Ortiz and Tye, 2014; Sun et al., 2019). Using a reporter for a glutamate transporter (vGlut2), we were able to show that the c-Fos induction occurs robustly, although certainly not exclusively, in a glutamatergic cell population in the BLA (**Fig. 7**).

Prior work has suggested that the microcircuit of the basolateral amygdala (BLA) to mPFC contributes to ASD-like behavior in mice. Felix-Ortiz and Tye (2014) used optogenetic approaches to either activate or inhibit BLA inputs to the mPFC during behavioral assays, and showed that the excited BLA/mPFC microcircuit reduced social interaction whereas suppression favoured social behavior. Sun and colleagues (2019) demonstrated that the glutamatergic neurons from the BLA preferentially projected to GABAergic neurons in the mPFC. Both their optogenetic and chemogenetic stimulations of this pathway resulted in reduced mPFC activity and stereotypic behavior. The ventral hippocampus (vHp) is the second downstream target. When the glutamatergic projection from the BLA to the vHp mice was activated, social interaction was reduced while the self-grooming was increased. This glutamatergic BLA/vHp pathway was further confirmed by the finding that glutamate receptor antagonists delivered to the vHPC attenuated optogenetic-evoked behavioral changes (Felix-Ortiz and Tye, 2014). Therefore, we speculate that the DLaN driven activation of the glutamatergic cell population in the BLA (**Table 7**) is what drives the grooming behavior observed in the present study.

With rapidly growing studies reporting the effects of lights on not only circadian rhythms but also the brain health (Stevenson et al., 2015; Lunn et al., 2017; Bumgarner and Nelson, 2021), the consequences of worldwide prevalence of DLaN should be taken into concern of disease management seriously, especially to those patients with neurodevelopmental disorders and neurodegenerative diseases who are vulnerable to environmental perturbations. Many people send the majority of time inside under artificial lighting, their exposure to DLaN needs to be carefully considered. Inappropriate light at night is common in hospitals, long-term care centers, and even our homes (Fournier and Wirz-Justice, 2010; Burgess and Molina, 2014; Osibona et al., 2021; Xiao et al., 2021). Thus, nightly light pollution is a very common environmental disruption to our circadian rhythms all over the world, and this present study shows that the undesired outcomes of DLaN on sleep/wake cycles and autistic behavior can be lessened by tailoring the spectral content of light sources to minimize melanopic stimulation.

## Abbreviations

ANOVA: analysis of variance
ASD: autism spectrum disorders
DD: constant darkness
*Cntnap2* KO: contactin associated protein-like 2 knock out
DLaN: dim light at night
ipRGCs: intrinsically photosensitive retinal ganglion cells
LD: light:dark
SEM: standard error of the mean
SCN: suprachiasmatic nucleus
λ: wavelength
WT: wild-type
ZT: Zeitgeber Time

## Acknowledgments

We would like to acknowledge UCLA students Kyle Nguyen-Ngo and Kelly Yun who helped with the scoring of some of the fluorescence staining. Dr. Kathy Tamai provided editorial assistance. Core equipment used in this study was supported by the National Institute of Child Health Development under award number: 5U54HD087101 (PIs Bookheimer, Kornblum). HBW was supported by a government scholarship for study abroad provided by the Taiwan Ministry of Education (#1100090595).

