## supplemental figures for "Long wavelength light reduces the negative consequences of dim light at night in the *Cntnap2* mouse model of autism"

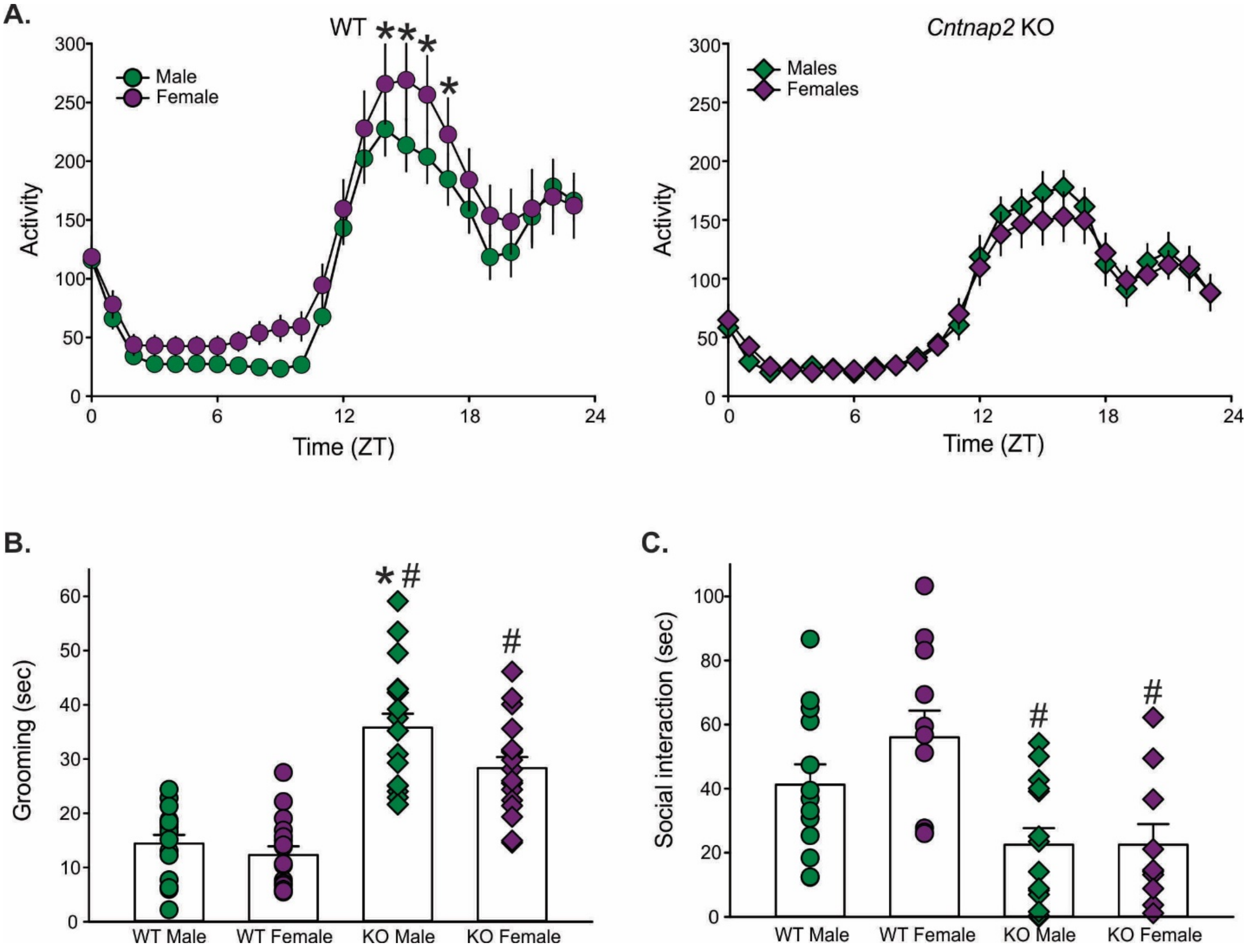


**S. Fig. 1: Sex differences between the WT and *Cntnap2* KO mice under baseline conditions (LD).** **(A)** Waveforms of daily rhythms in cage activity in WT (circles) and *Cntnap2* KO (diamonds) male (green) and female (purple) mice. The activity waveform (1 hr bins) of each group was analyzed using a Two-way ANOVA for repeated measures with time and sex as factors followed by the Holm-Sidak multiple comparisons test. The WT mice exhibited significant effects of time ($F_{(23,551)}$ = 31.64, $P$ < 0.001) and sex ($F_{(1,551)}$ = 20.99, $P$ < 0.001) but no interaction between the two factors was identified ($F_{(23,551)}$ = 0.55, $P$ = 0.959). The *Cntnap2* KO exhibited significant effects of time ($F_{(23,575)}$ = 38.92, $P$ < 0.001) but not of sex ($F_{(1,575)}$ = 0.93, $P$ = 0.336) as well as no significant interaction between the two factors ($F_{(23,575)}$ = 0.37, $P$ = 0.997). Asterisks indicate significant ($P$ = 0.959) differences between the 1 hr bins as measured by the Holm-Sidak test for multiple comparisons. (**B**) Time spent grooming by WT and *Cntnap2* KO male and female mice. Grooming behavior was assessed in a novel arena at ZT 18. (**C**) Social behavior was assessed by measuring the time each mouse spent actively interacting with a novel mouse, matched by age, sex and genotype, at ZT 18. WT and *Cntnap2* KO male and female mice. Data in panels **B** and **C** were analyzed using a two-way ANOVA with genotype and sex as factors followed by the Holm-Sidak multiple comparisons test and are reported in **Table 4**. Histograms show the means ± SEM with overlaid the values from individual animals. Significant (P < 0.05) effects of treatment or genotype are indicated with an asterisk or a crosshatch, respectively.

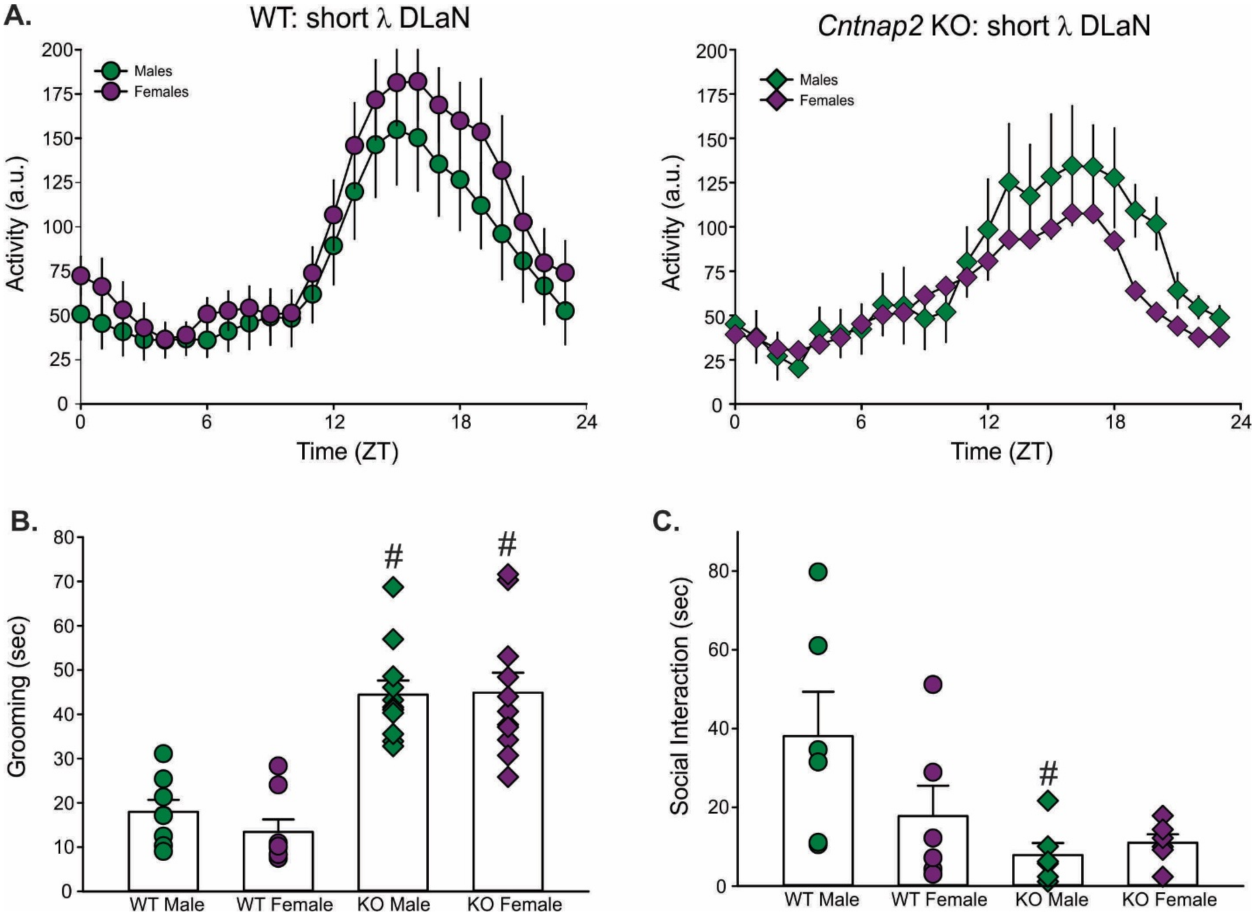


**S. Fig. 2: Lack of sex differences between WT and *Cntnap2* KO mice exposed to DLaN with a short λ enriched light.** WT (circles) and *Cntnap2* KO (diamonds) male (green) and female (purple) mice were exposed to the short λ enriched DLaN for two weeks. **A)** Waveforms of daily rhythms in cage activity. The activity waveform (1 hr bins) of each treatment group was analyzed using a two-way ANOVA for repeated measures with sex and time as factors followed by the Holm-Sidak multiple comparisons test. Both genotypes exhibited significant effects of time (WT: $F_{(23,287)} = 13.36$, $P < 0.001$; *Cntnap2* KO: $F_{(23,287)} = 5.32$, $P < 0.001$) and sex (WT: $F_{(1,287)} = 12.61$, $P < 0.001$; *Cntnap2* KO: $F_{(1,287)} = 5.99$, $P = 0.015$), but no interaction between the two factors was identified (WT: $F_{(23,287)} = 0.21$, $P = 1.000$; *Cntnap2* KO: $F_{(23,287)} = 0.42$, $P = 0.992$). (**B**) Time spent grooming by WT and *Cntnap2* KO male and female mice. Grooming behavior was assessed in a novel arena at ZT 18. (**C**) Social interactions were assessed by measuring the time each mouse spent actively interacting with a novel stranger mouse at ZT 18. Data in panels **B** and **C** were analyzed using a two-way ANOVA with genotype and sex as factors followed by the Holm-Sidak multiple comparisons test and are reported in **Table 5**. Histograms show the means ± SEM with overlaid the values from individual animals. Significant ($P < 0.05$) effects of genotype are indicated with a crosshatch.
